# Enhancing the biological relevance of Gene Co-expression Networks: A plant mitochondrial case study

**DOI:** 10.1101/682492

**Authors:** Simon R. Law, Therese G. Kellgren, Rafael Björk, Patrik Ryden, Olivier Keech

**Author notes:** Author for correspondence: Olivier Keech, Department of Plant Physiology, Umeå Plant Science Centre, Umeå University, S-90187 Umeå, Sweden.

## Abstract

Gene Co-expression Networks (GCNs) are obtained by a variety of mathematical of models commonly derived on data sampled from diverse developmental processes, tissue types, pathologies, mutant backgrounds, and stress conditions. These networks aim to identify genes with similar expression dynamics, but are prone to introduce false-positive and -negative relations, especially in the instance of large and highly complex datasets. With the aim of optimizing the relevance of edges in GCNs and enhancing global biological insight, we propose a novel approach that involves a data-centering step performed simultaneously per gene and per sub-experiment, called *centralisation within sub-experiments* (CSE).

Using a gene set encoding for the plant mitochondrial proteome as a case study, our results show that CSE-based GCNs had significantly more edges within the majority of the considered functional sub-networks, such as the mitochondrial electron transport chain and its sub-complexes, than GCNs not using CSE; thus demonstrating that the CSE-based GCNs are efficient at predicting those canonical functions and associated pathways, also referred to as the “core network”. Furthermore, we show that CSE, in conjunction with conventional correlation analyses can be used to fine-tune the prediction of the function for uncharacterised genes; while in combination with analyses based on non-centralised data can augment those conventional stress analyses with the innate connections underpinning the dynamic system examined.

Therefore, CSE appears as an alternative method to conventional batch correction approaches. The method is easy to implement into a pre-existing GCN analysis pipeline and can provide accentuated biological relevance to conventional GCNs by allowing users to delineate a “core” gene network.

**Author Summary:** Gene Co-expression networks (GCNs) are the product of a variety of mathematical models that identify causal relationships in gene expression dynamics, but are prone to the misdiagnoses of false-positives and -negatives, especially in the instance of large and highly complex datasets. In light of the burgeoning output of next generation sequencing projects performed on any species, under different developmental or clinical conditions, the statistical power and complexity of these networks will undoubtedly increase, while their biological relevance will be fiercely challenged. Here, we propose a novel approach to primarily generate a “core” GCN with augmented biological relevance. Our method, which involves data-centering steps and thus effectively removes all primary treatment / tissue /patient effects, is simple to employ and can be easily implemented into pre-existing GCN analysis pipelines. The gained biological relevance of such an approach was validated using a subcellular gene set encoding for the plant mitochondrial proteome, and by applying numerous steps to challenge its application.

## Introduction

Over the last two decades, the exponential growth of available transcriptome data in an increasing number of species has given rise to the establishment of a multitude of gene co-expression networks (GCNs). By constructing these networks on data sampled from diverse developmental processes, tissue types, pathologies, mutant backgrounds or stress conditions, researchers can better comprehend the physiological and molecular pathways that underpin complex biological systems (Carrera et al., 2009; Emmert-Streib et al. 2014; Castro et al. 2019). These networks are reliant on mathematical models to identify causal relationships in gene expression dynamics, and although there are many different permutations of these models, the most prevalent are those based on conventional correlation approaches such as Pearson correlation coefficient or Spearman’s rank correlation coefficient.

These conventional correlation methods have been demonstrably successful at identifying cohorts of strongly co-expressed genes, and thus extensively used to generate GCNs; however, these methods also have their drawbacks. This is especially apparent in large and complex datasets where a large fraction of the predicted correlations are expected to be statistically significant, and causal gene-to-gene connections are obscured by the over-whelming presence of false -positives and -negatives. Non-causal relationships can arise from indirect connections with other gene products and from non-biological sources such as influences stemming from experimental design. Partial correlation is a standard approach used to attenuate non-causal relationships generated by the influence of other genes. One such approach, Gaussian Graphical Modelling, is commonly used as it allows researchers to interrogate the direct association between two genes, independent of the effects of surrounding genes present in the dataset. A number of thorough Gaussian Graphical Modelling studies in the model plant species *Arabidopsis thaliana* (Arabidopsis) have demonstrated the statistical power of this technique, and generated GCNs of select pathways and on a genome-wide scale (Wille et al., 2004; Ma et al., 2007; Ma et al., 2015). Another useful step for analysing complex datasets encompassing a wide range of tissues, developmental stages and stresses, is the use of batch-effect removal approaches. Conventional batch-effect removal approaches effectively eliminate the systematic, technical errors inherent to multi-experiment comparisons (Chen et al. 2011; Nygaard et al. 2016). However, GCNs obtained utilizing partial correlation and batch-effect removal approaches will not reduce non-causal relationships caused by unquantifiable factors, e.g. treatment/tissue effects between samples. Hence, there is currently a lack of methodology to robustly derive informative GCNs from complex datasets generated by multiple experiments.

Although these GCNs are not an end on to themselves, they can be utilised in a number of innovative ways to reveal evidence of functions for otherwise uncharacterised gene products, identify novel protein localisation, and to better describe complex biological pathways which can react fluidly to different stresses/developmental processes. In the field of plant science, GCN inferences have been utilised to great effect, particularly in the model species Arabidopsis, both to construct genome-spanning global networks and biological sub-pathways (Liesecke et al., 2018). Yet, beyond its statistical value, the biological relevance of an edge between two nodes in such networks can rightfully be questioned. Furthermore, validation of GCNs can be challenging, as there are a limited number of gene-to-gene relationships demonstrated to be positive (i.e. a causal relationship) or negative (i.e. a non-causal relationship) (Qian and Dougherty, 2013; Chai et al. 2014; Banf and Rhee, 2017). An alternative is to base the validation (i.e. the biological relevance of the output) on functional proximity and functional categories, arguing that the fraction of causal relationships should be relatively high within sets of genes encoding proteins that comprise the same protein complex or are involved in the same metabolic pathway.

With the aim of optimizing the biological relevance of edges in GCNs and enhancing global biological insight, we challenged different methodologies in the generation of these networks by using, for biological validation, a subset of nuclear genes encoding plant mitochondrial-targeted proteins (As defined in Chrobok et al., 2016). To achieve this, we applied a novel pre-processing step that we called *centralisation within sub-experiments* (CSE), which removes batch-effects and reduces the impact of confounding effects of treatment-induced and tissue-specific responses. In contrast to conventional batch-effect removal approaches, the centralisation step is applied to datasets where the observations are biological replicates derived under the same experimental conditions. Hence, CSE removes treatment-induced or tissue-specific effects, and technical bias introduced by variability between experiments. Here, GCN approaches based on either Pearson or partial correlation using CSE were compared to corresponding approaches in the absence of the CSE step. The biological validation was conducted by categorizing a subset of genes encoding for plant mitochondrial proteins with respect to expression patterns, functional proximity, and functional categories. CSE combined with GCN (utilizing Pearson correlation) provided the optimum balance of ease of data processing vs. the utility of the output. Consequently, a mitochondrial network based on CSE Pearson correlation was selected for further downstream applications of the method.

## Results

To gain clarity, this results section has been divided in 3 parts: Methodology, Validation and Application.

## Methodology

### Definition of the problem

We consider a problem where we have gene expression data from a large number of diverse experiments, e.g. experiments from different tissues, conditions and developmental stages. The objective is to predict the edges of an undirected graph with *n* nodes (i.e. genes), where an edge represents the most pronounced co-expression between a pair of genes. Often, the level of co-expression between genes will be context-dependent, e.g. tissue, growth condition or developmental stage. Here we are primarily interested in detecting the core network, i.e. to estimate the co-expression between genes that are prominent in the majority of the considered sub-experiments. A sub-experiment is defined as a set of assays derived under “identical settings”, i.e. the assays within the sub-experiment can be treated as biological replicates. We thus propose a pre-processing step (CSE) that enables prediction of the core network.

### Centralisation within sub-experiments

We consider normalized gene-expression data from *s* sub-experiments i.e.

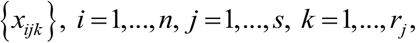

where *x*_*ijk*_ denotes the normalized gene-expression for gene *i* observed on the *k*^th^ biological replicate in sub-experiment *j*. CSE is a simple pre-processing step whereby mean-centralization within sub-experiments is applied to each gene separately, i.e. the CSE-processed expressions are obtained as:

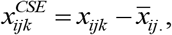

where 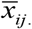 denote the mean-expression of gene *i* in the *j*^th^ sub-experiment, *i* = *1*,…,*n*, *j* = *1*,…,*s*, *k* = *1*,…,*r*_*j*_.

It should be noted that the mean value of the centralised data within a sub-experiment will always be zero. The idea behind CSE is to avoid pronounced correlations driven by differences between the sub-experiments. For example, a stress may induce gene expression in genes that are expressed in “independent” pathways resulting in false positive and false negative predictions (Supp. Fig. 1).

### Construction of Gene Co-expression Networks

GCNs can be constructed in various way, but we focused on some commonly used conventional approaches to assess the effect of CSE application. The GCNs were constructed in a three-step procedure: (i) the pre-processed dataset was either centralised using CSE (C) or not centralised (NC), (ii) pairwise correlations were calculated using either Pearson correlation (PeC) or partial correlation (PaC), and (iii) the *sign matrix* (i.e. an adjacency matrix whose entries are either 1, 0 or −1) was constructed by controlling the fraction ω of edges at a desired level, i.e. controlling the sparcity at level ω. The network was defined by the output of the precision matrix; where a “one” represents an edge corresponding to a level of co-expression between genes that satisfies a given cut-off. In this study, four different principal networks were evaluated: combining CSE and Pearson correlation (CPeC), CSE and partial correlation (CPaC), and Pearson and partial correlation applied in the absence of CSE (NCPeC and NCPaC, respectively). The sparsity of all GCNs was controlled at ω=0.005 and the Walktrap community detection algorithm (Pons and Latapy, 2005) was used to identify communities in the predicted GCN based on Pearson correlation, see Method for further details.

It should be noted that the objective here was not to predict all edges in the core network, but to predict the most pronounced edges, which justifies the use of an arbitrary chosen threshold. Moreover, having the same sparsity in all predicted networks simplified the evaluation as described below.

Applying the conceptual reasoning outlined above on a network using simulated data demonstrated that CPaC removes non-causal edges arising from the influence of other genes and non-causal edges caused by external factors (Supp. Fig. 1). Similar results were obtained for CPeC, with the exception that a few false, but relatively weak, edges appeared. The network utilising non-centralised data and Pearson correlation, arguably a fairly standard approach, resulted in dense networks with several false positives. Due to computational constraints, partial correlation approaches may not be suitable for constructing GCNs when the number of genes is much larger than the number of experiments, see Discussion.

### Evaluation of Gene Co-expression Networks

We consider a core network *C*, with *n* nodes and *k* edges, where the edges corresponds to the fraction ω of the strongest co-expression correlation. A sub-network *A ⊂ C*, with *n*_*A*_ nodes and *k_A_* edges is said to be pronounced if *k*_*A*_ is larger than the expected number of edges in a randomly selected sub-network with *n*_*A*_ nodes, i.e.

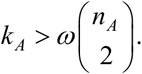

The network *C* is commonly unknown, but it may still be possible to identify several pronounced sub-networks, e.g. by considering physical or functional proximity, see Methods - Preparing elements of the mitochondrial working model (iii-iv).

We propose that the relative performance of predicted GCNs, all with the same sparsity ω, can be evaluated based on the observed number of edges within pronounced sub-networks. In short, we argue that the more observed edges (the lower p-values) within pronounced sub-networks, the better the predicted networks are, see Methods for further details. With that being said, there is a risk to overestimate the number of edges within the pronounced sub-networks resulting in an incorrect ranking of the considered networks, however, this risk decreases as the number of pronounced sub-networks is increased.

### Validation

For this study, we chose the plant mitochondrion as a focal point for 3 main reasons: (i) assessing the biological relevance of our findings became much easier due to our pre-existing knowledge on the plant mitochondrial metabolism, (ii) the number of genes to work with is low (ca. 1000 nuclear genes encoding for mitochondrial-targeted proteins), hence easing the application of partial correlation methods, and (iii) the interest in mitochondrial biology is undoubtable as this organelle is recognised as a central energetic, signalling, and stress response hub in (almost) all eukaryotic cells.

### The effect of tissue type on Gene Co-Expression Networks

Visualisation of the four GCNs generated using Cytoscape (Organic layout; Shannon et al., 2003) revealed networks that shared strong similarities in structure depending on whether CSE was applied or not (Figure 2; Supplemental Table 2). Those networks based on non-centralised data displayed two distinct primary clusters of nodes (Figure 2A and B), while those based on centralised data were more integrated (Figure 2C and D). To uncover the source of these distinct clusters in the non-centralised data, we returned to the original data from the AtGenExpress expression atlas, and defined each gene as presenting dominant expression in either below-ground tissues (e.g. roots) or above-ground photosynthetic tissues (e.g. shoots and leaves), see Methods for details. Using these definitions, nodes (genes) from the Cytoscape-generated networks were coloured based on their classification as either below-ground dominant (brown), above-ground dominant (green) or dominance in neither tissue (yellow) (Figure 2). This rapidly demonstrated the strong influence tissue-of-origin has over the resulting co-expression networks, and the efficacy of CSE in resolving this. Notably, in addition to the increased integration of genes with different tissue-dominances, the number of nodes present (thus, the number of nodes with at least one edge to another node) in the four networks was significantly larger following CSE. Furthermore, the distribution of genes with tissue-dominance established an increased inclusion of genes with no tissue dominance (Neither), which brought these networks closer to the native distribution of tissue of origin dominance observed in the total set. This suggests that by removing external biases, CSE of data could introduce a wider cross-section of genes into a GCN and thus reveals novel interactions.

**Figure 1.**
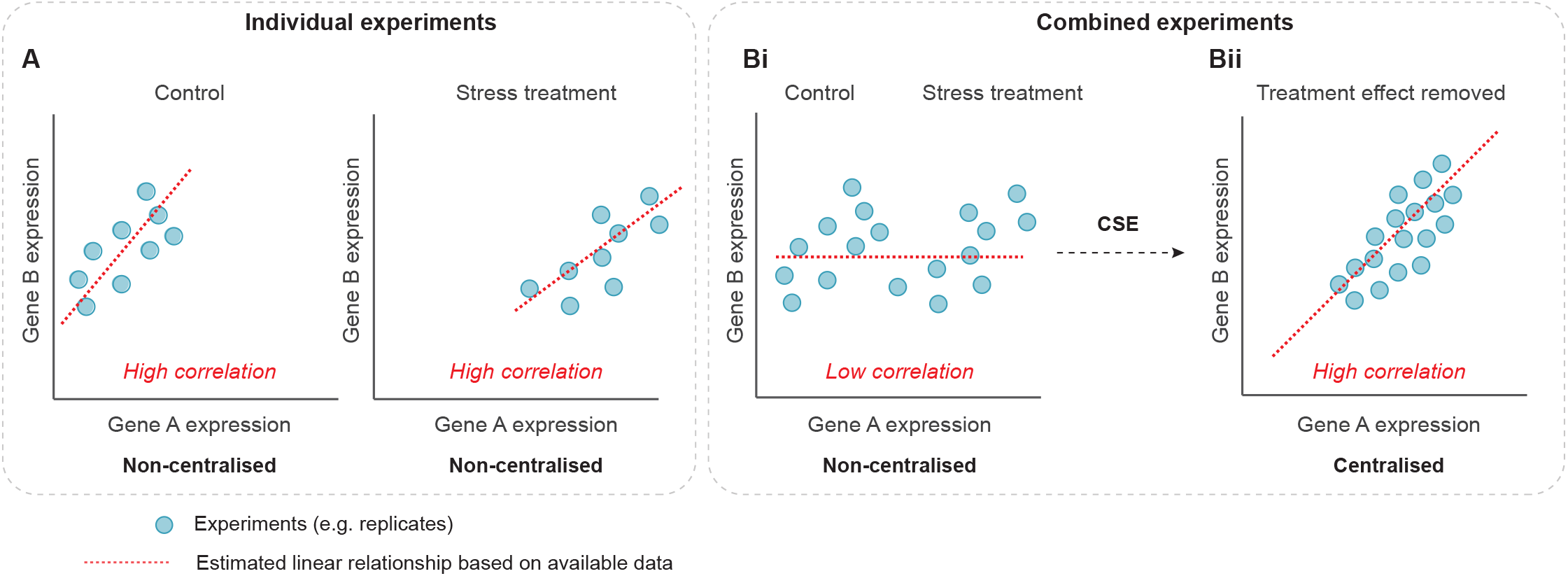
Schematic illustrating the utility of centralisation when comparing genes from a diverse background of treatments. **(A)** Conventional correlation analysis of two genes (Gene A and Gene B) under control conditions reveals a high positive correlation. Coresponding correlation analysis of the same two genes in response to a stress treatment again reveals a high positive correlation. **(Bi)** When both the control and stress experiments are combined, conventional correlation analysis results in a low level of correlation (false negative). **(Bii)** By carrying out centralisation within sub-experiments (CSE), the mean effect between replicates is removed, and subsequent conventional correlation analysis now reveals the “core” high correlation between Gene A and Gene B.

**Figure 2.**
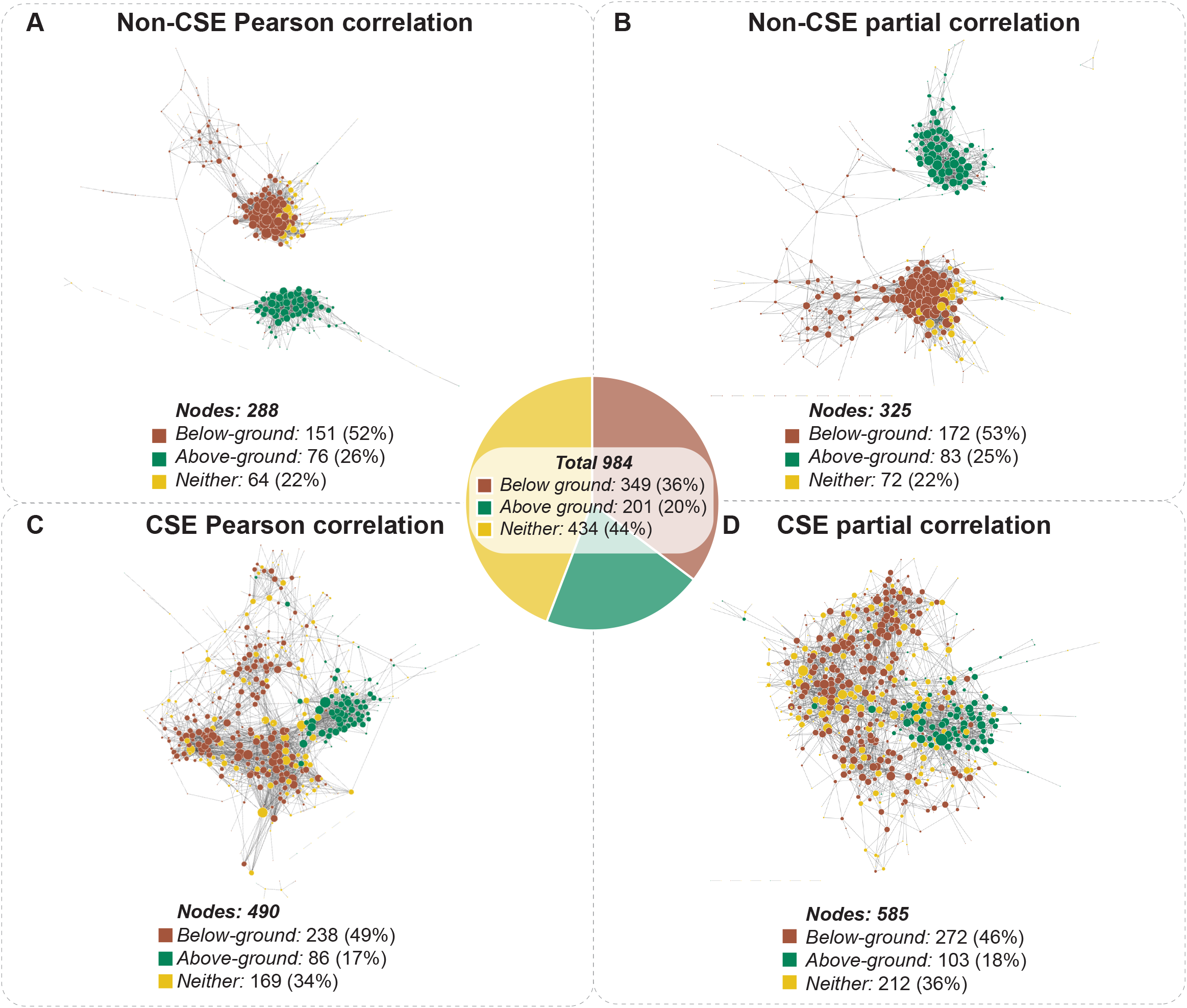
Visualisation of the mitochondrial network using four different pre-processing and correlation approaches. A manually curated mitochondrial gene list was cross-referenced with the AtGenExpress Expression Atlas spanning different tissues, developmental stages, and stresses (Schmid et al., 2005, Kilian et al., 2007 and Goda et al., 2008). This data was either subject to CSE or left unprocessed, prior to correlation analysis using either Pearson correlation or partial correlation. Each of the four resulting networks was visualised using Cytoscape. For each network, only nodes with at least one edge to another node were included. Each node (gene) was coloured based on their classification as either below-ground dominant (brown), above-ground dominant (green) or dominance in neither tissue (yellow). The diameter of each node is proportional to the number of edges it has to a neighbouring node. **(A)** Network of non-CSE Pearson correlation (**B)** Network of non-CSE Partial correlation **(C)** Network of CSE Pearson correlation (**D)** Network of CSE Partial correlation.

### Assessing interactions based on functional proximity

Our first approach at challenging the four different co-expression networks was to examine the resulting distribution of edges upon a small isolated subset of the mitochondrial network, encoding components of the mitochondrial electron transport chain (mETC). The mETC is central to the bioenergetic function of mitochondria and the array of genes that comprise its five complexes have been demonstrated to be expressed at relatively stable levels in a variety of tissue types and developmental stages (Lee et al., 2011). A comparison between the mETC set isolated from the four networks revealed a significantly higher number of edges (derived from connections within and between the five complexes) in the networks based on centralised data, while the influence of Partial correlation vs. Pearson correlation was comparatively small (Figure 3A). As the same sparsity is applied to all four approaches, the total number of edges in the entire network is held consistent between them; thus, the enrichment of edges within the mETC observed here represents a valuable indication of putative biological interaction. Our next step was to assess the distribution of edges within a single complex of the mETC. The NADH dehydrogenase, commonly known as Complex I, is composed of three domains: the peripheral arm domain (PAD), the membrane arm domain (MAD), and the carbonic anhydrase domain (CAD) (Klodmann et al. 2010). Each domain in turn is composed of an assembly of proteins that carry out highly specialised functions, and thus proved ideal to assess the relevance of the distribution of edges between the four different approaches (Methods; Figure 3B). Similar to the distribution of edges for the entire mETC, there were far more edges between the nodes of Complex I in centralised than in non-centralised data, while the difference in the number of edges between networks based on Pearson or Partial correlation is negligible (Supplemental Table 3). In almost all cases, when comparing the number of edges expected to occur by chance between the genes of the three domains with the actual observed edges, this enrichment was found to be highly significant (Figure 3B). When this examination was expanded to look at the distribution of edges within and between all five complexes of the mETC, a similar enrichment of significant interactions was observed in the centralised data, but not in the non-centralised data (Figure 3C). Interestingly, when the distribution of edges between individual complexes and either (i) pooled complexes of the mETC, or (ii) the rest of mitochondrial set (total mitochondrial set, excluding the mETC), the networks based on non-centralised data showed relatively poor correlations with the pooled mETC and even weaker connections with the non-mETC components (Figure 3D). In contrast, the centralised data showed significantly (P<0.001) strong connections between the individual complexes and the pooled mETC, with weaker connections to the non-mETC components. One important exception to this was the significant (P<0.001 in CPaC, and P<0.01 in CPeC) connection observed between Complex II and the non-mETC components. Notably, Complex II (also called Succinate Dehydrogenase) lies at the confluence of two essential bioenergetic functions of the mitochondrion: the mETC and the TCA cycle. As such, it is particularly notable that the centralised data identified Complex II as having significant interaction with non-mETC components. Examination of the composition of edges between Complex II and these non-mETC genes revealed that they were indeed significantly (P<0.0001) enriched in components of the TCA cycle. Taken together, these observations strongly support that CSE of data prior to correlation analysis can reveal gene-to-gene interactions indicative of highly valuable biological relationships such as association to shared protein domains or consecutive enzymes in a metabolic pathway.

**Figure 3.**
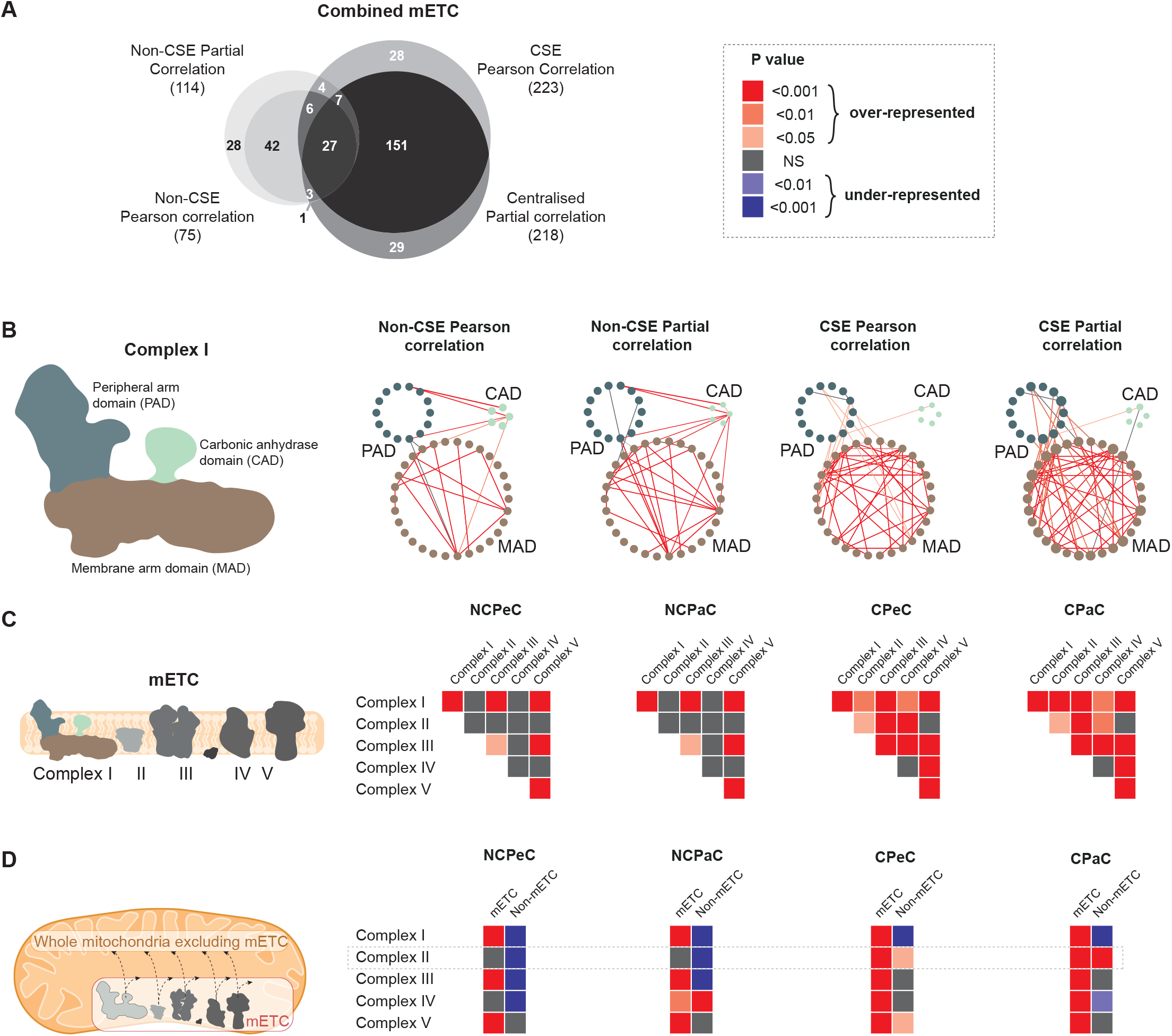
Comparative analysis of four different correlation methods in defining interactions based on functional proximity. The following gene subsets of the mitochondrial electron transport chain were analysed using non-CSE Pearson correlation (NCPeC), non-CSE Partial correlation (NCPaC), CSE Pearson correlation (CPeC) and CSE Partial correlation (CPaC). P values were calculated for the probability associated with the expected vs. observed number of edges and a colour-grading scheme of the resulting P values applied. **(A)** A Venn diagram illustrating the overlap of connections between the complexes of the mitochondrial electron transport chain (mETC), when analysed using the four different correlation methods. **(B)** The significance of the edges between the three domains of Complex I. **(C)** The significance of the edges within a given complex or between the different complexes of the ETC. **(D)** Between the individual complexes of the mETC vs. the unified mETC or the rest of the mitochondrial set excluding the mETC.

### Assessing interactions based on connectivity within and between mitochondrial functional categories

Using the newly updated functional annotations established for the MapMan platform (MapMan X4 Release 1.0, 2018; Usadel et al., 2009), each gene of the mitochondrial set was assigned to one of 29 functional categories. By grouping genes belonging to the same functional categories, we were able to measure the number of edges between genes *within* a functional category, versus those between *different* but interrelated functional categories (Figure 4). In brief, when CSE had been carried out (Figure 4C and D), the number of significant edges between genes within the same category is much higher (ca. doubled) than is observed when the data is non-centralised (Figure 4A and B). Additionally, in the two centralised datasets, the number of significant edges between *different* functional categories also increases, when compared to their non-centralised counterparts. These inter-category edges were often highly biologically relevant: for example a significant (P<0.0001) edge was observed between *nucleotide metabolism* and *protein biosynthesis* in each of the four methodologies (Figure 4A-D), which is hardly surprising given their canonic interconnectivity. In contrast, some connections were only observed in the case of the centralised datasets (Figure 4A-B), such as the significant (P<0.0001) edges between *cellular respiration* and *carbohydrate* and *lipid metabolism*, as well as the connection between *protein biosynthesis and protein translocation*. For these processes to operate efficiently, a high level of coordination in the regulation of the genes involved is required, which supports these additional inter-category edges. In summary, the known biological pathways strongly corroborate the input from the centralized co-expression data generated with our mitochondrial dataset and undoubtedly strengthen its consideration for future analyses. Following these validation steps, the negligible difference in results between centralised Pearson correlation and partial correlation, contrasted with computational demands associated with the latter, led us to progress with the subsequent application using centralised Pearson correlation (CPeC).

**Figure 4.**
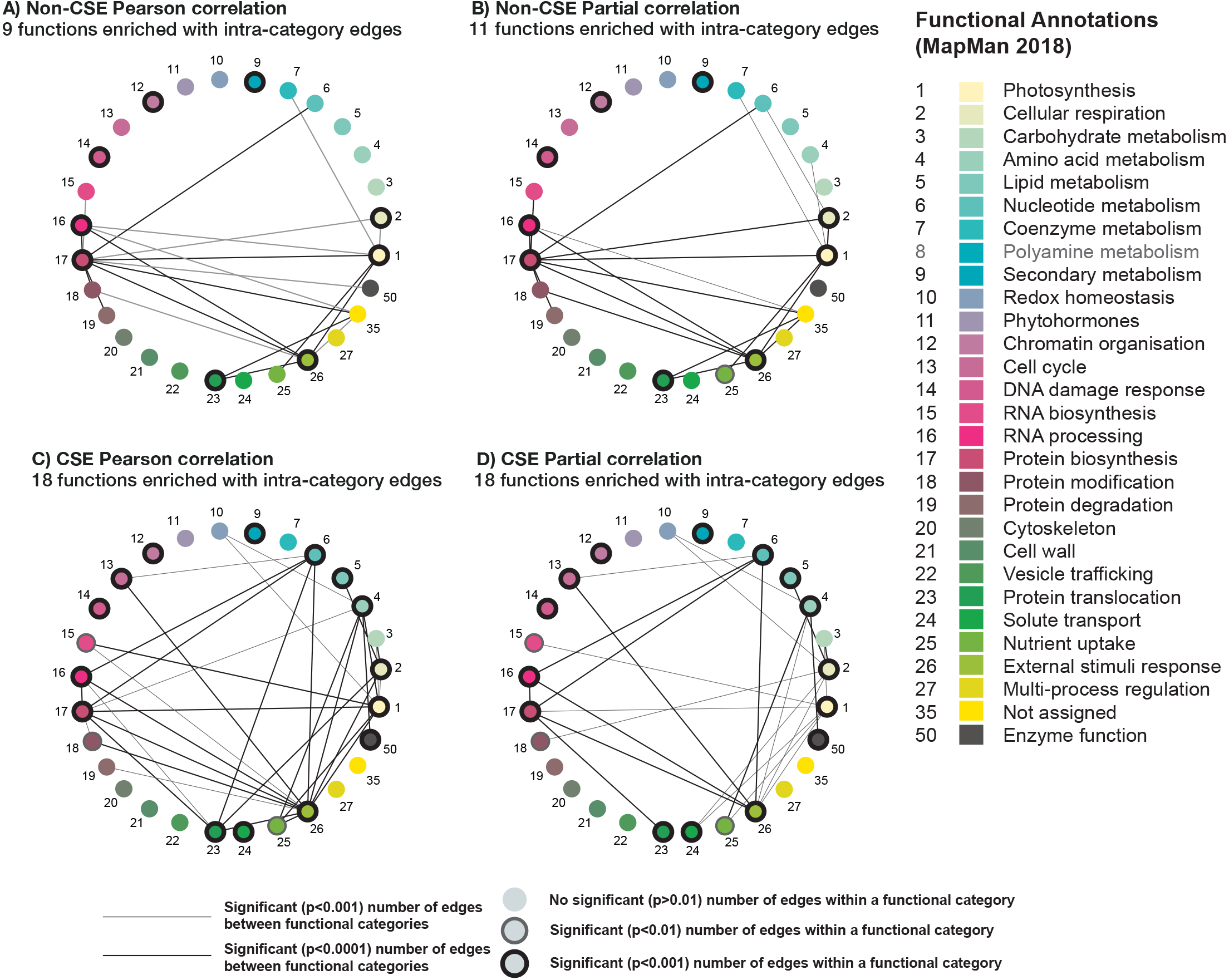
Comparative analysis of four different correlation methods based on connectivity between different functional categories in mitochondria. Using newly updated MapMan annotations (MapMan X4 Release 1.0, 2018; Usadel et al., 2009), the mitochondrial set was subdivided into 29 different functional categories. Only functional categories with at least one significant connection to another category are displayed for each method. Nodes with a black outline indicate functional categories with significant intra-connectivity, nodes lacking an outline indicates functional categories that do not have significant intra-connectivity. **(A)** Non-CSE Pearson correlation (NCPeC), **(B)** Non-CSE Partial correlation (NCPaC), **(C)** CSE Pearson correlation (CPeC), and **(D)** CSE partial correlation (CPaC).

## Application

### Using the network to predict the function of an uncharacterised mitochondrial gene

The functional annotations applied to the genes comprising the mitochondrial network (introduced above) encompassed a subset of mitochondrial genes that at the time of the publication of the MapMan hierarchical set of functional categories (BINs; MapMan X4 Release 1.0, 2018), encoded proteins with no assigned functions (NAFs; Functional Category 35). This provided an ideal target group that we could systematically interrogate, in a “guilt by association manner”, to determine if their relationship to other genes of known functions could support their putative function. A subsequent mitochondrial network was established, which comprised 111 NAF genes and 257 mitochondrial genes encoding proteins with known functions that had at least one edge to a NAF gene (Figure 5A; Supplemental Table 4). The NAF genes were then arranged in descending order based on those with the greatest number of edges to genes with known functions. We then selected the top 5 NAF genes and identified the genes they interacted with. Next, the distribution of their associated functional annotations was determined and assessed to see if they were enriched in a particular function (Figure 5B).

**Figure 5.**
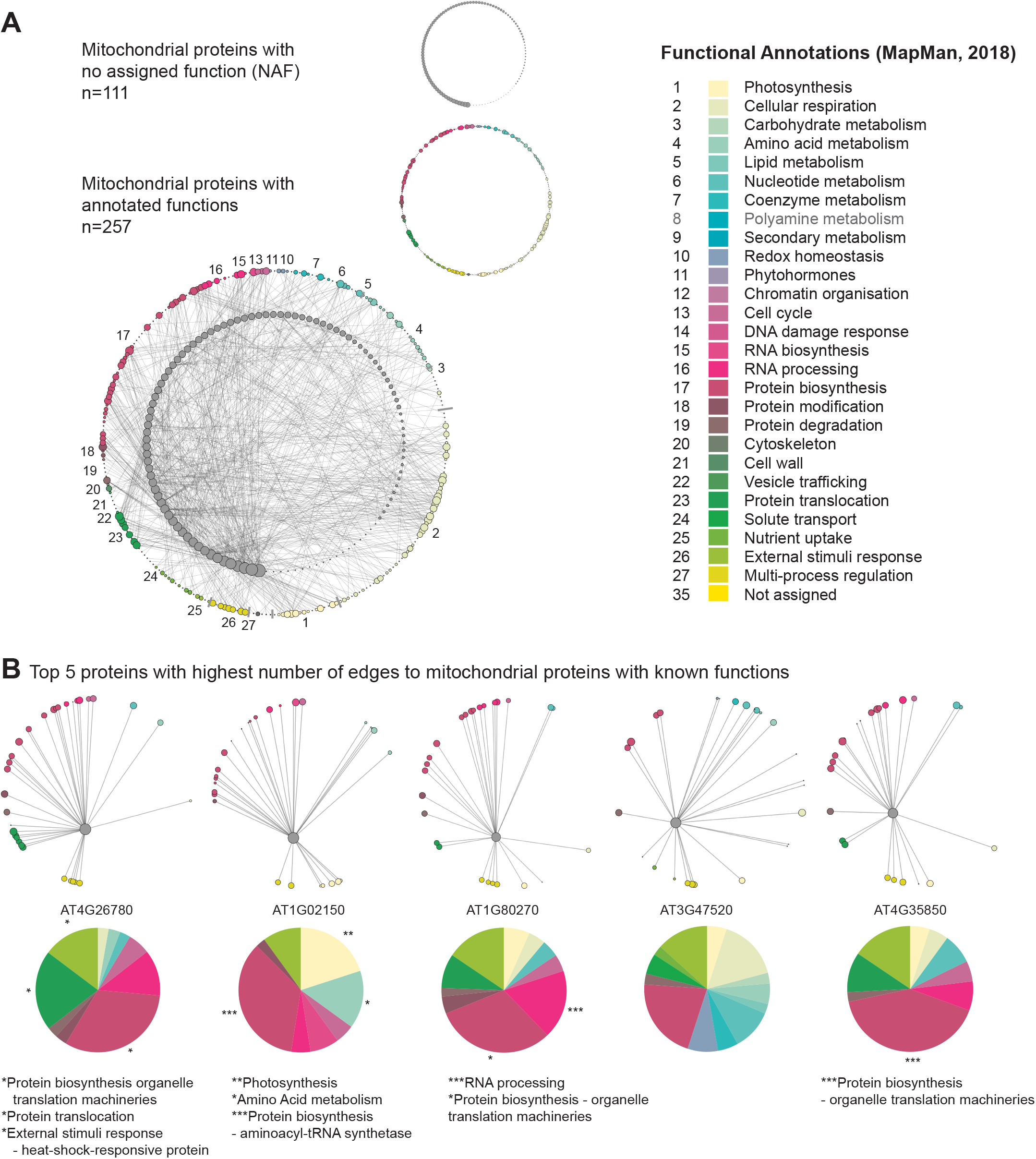
Identification of candidate functions for mitochondrial proteins with unknown functions. Pearson correlation was carried out on centralised data spanning the mitochondrial set, over 370 unique conditions comprising the AtGenExpress Expression Atlas. Out of this list, a sub-population of genes was established which had unknown functional annotations. This sub-population was then analysed to identify significant interactions with mitochondrial proteins with known functions, resulting in a suite of 109 mitochondrial proteins with unknown functions. By annotating the functional categories of the known mitochondrial genes, putative functional relationships can be assigned to these as yet uncharacterised proteins. **(A)** Network representation of the interactions between 109 mitochondrial proteins with no annotated functions and 248 mitochondrial proteins with known functions. **(B)** The five proteins with no annotated functions displaying the highest number of edges to the mitochondrial set are shown, with a functional breakdown of the distribution of edges. Significant over-representation of a given functional category has been marked with the following: p<0.05 = *; p<0.01 = **; p<0.001 = ***.

The top 5 NAF genes displayed significant over-representations with a range of different functional categories. The NAF with the greatest number of connections with genes of known function, AT4G26780, had a significant enrichment of edges with (i) protein biosynthesis - organelle translation machineries (P<0.05), (ii) protein translocation - TOM translocation and TIM insertion systems (P<0.05), and (iii) external stimuli response - heat-shock-responsive protein (P<0.05). Interestingly, this protein has been proposed to encode Mge2, which is one of two mitochondrial GrpE proteins in Arabidopsis. The remaining homologue, Mge1 serves as a co-chaperone alongside Hsp70, which together form a vital part of the presequence-assisted motor (PAM) complex that aids in the transport of precursor proteins through the TIM17:23 translocase (Hu et al., 2012; Ghifari et al., 2018). While Mge1 appears to have more constitutive house-keeping duties, Hu et al., (2012) demonstrated that Mge2 was specifically induced by heat and suggested that it could be required for mitochondrial protein import and folding during periods of heat stress, a hypothesis that appears to be supported by our GCN predictions. The second gene interrogated (AT1G02150), had a significant enrichment of edges with (i) photosynthesis functions (P<0.01), (ii) amino acid metabolism (P<0.05), and (iii) protein biosynthesis - aminoacyl-tRNA synthetase (P<0.001). At present, little is known about this protein, however, the Arabidopsis Information Portal (Araport) 11 classifies it as belonging to the tetratricopeptide repeat (TPR)-like superfamily (Cheng et al., 2017). TPR domains can be found in a diverse number of proteins, where they mediate protein-protein interactions; particularly in the formation of protein complexes. The strong significant (P<0.001) over-representation with aminoacyl-tRNA synthetase functions (and the weaker, though still significant over-representation of amino acid metabolism functions) observed here is particularly interesting, as there is evidence that TPR-containing proteins can act as interacting mediators and co-chaperones in the formation of aminoacyl-tRNA synthetases (Han et al., 2007; Kim et al., 2014); suggesting that this protein may have a role in assisting amino acid loading of tRNAs in Arabidopsis. The third gene interrogated (AT1G80270) had a significant enrichment of edges with (i) RNA processing (P<0.001) and (ii) protein biosynthesis - organelle translation machineries (P<0.05). Assessing the available literature, this protein has been reported as belonging to the pentatricopeptide (PPR) superfamily (Doniwa et al., 2010), which are predominately mitochondrial or plastid targeted proteins and have been demonstrated to have a diverse array of roles associated with RNA metabolism, such as RNA editing, splicing, stability and translation (Barkan and Small, 2014). AT1G80270, known as PPR596, has been demonstrated to be involved in the C-to-U editing efficiency of ribosomal protein S3 (RPS3; AtMg00090), which is noteworthy as in our study, PPR596 was also significantly enriched in connections with organelle translation machinery functions (Doniwa et al., 2010). Regarding AT3G47520, despite the surprising lack of a proper annotation by Mapman, this gene had been characterized and encodes an isoform of the mitochondrial dehydrogenase (mMDH2; Tomaz et al., 2010; Lindén et al., 2016). Although no functional categories were enriched, the big proportion taken by the categories redox homeostasis, cellular respiration and protein biosynthesis strongly supports the physiological role of mMDH2. Finally, the protein encoded by AT4G35850 had a significant (p<0.001) enrichment of edges with protein biosynthesis - organelle translation machineries (large and small mitoribosome subunit) functions. Very little is known about this protein, but it has been classified as belonging to the PPR superfamily by Araport11, and could thus have a similar role to that of PPR596; as an editing factor associated with the correct processing of transcripts encoding mitoribosomal subunits, or be associated with ribosomes in other ways described in the literature; such as maintaining the stability of assembled mito-ribosomes following translation (Schmitz-Linneweber and Small, 2008); or promoting translational initiation by selectively recruiting mitoribosomes to the start codon of their target transcripts (Manavski et al., 2012; Haïli et al., 2016). Taken together, these findings suggest that guilt by association-style analysis of networks founded on data subjected to CSE offers an attractive first step in the process of characterising genes where little is known about them.

### Synergy of centralisation approaches in the analysis of plant stress

In the field of transcriptomics, the application of conventional co-expression networks has proven a highly powerful approach in characterising stress responses in a diversity of organisms. In this study, we have demonstrated that CSE of data prior to correlation analysis effectively identifies the innate relationship between genes, and thus delineates a “core gene-network”. However, as previously mentioned, a caveat of this approach is that it is predicated on the suppression of extraneous effects, such as stress, tissue, treatment, or genotype from a given dataset, which therefore prevents us from interrogating the impact of these outside influences on the dynamics of the co-expression network generated. On the other hand, quite often researchers must adjust different parameters (cut-offs, thresholds, etc.) to introduce enough genes to reposition the stress-responsive network in a wider biological context and gain understanding. Here we propose an alternative method, with a powerful reference tool that can augment conventional co-expression analyses. By clustering the CSE data of the entire AtGenExpress Expression Atlas using a Walktrap community detection algorithm (Pons and Latapy, 2005), we generated a hierarchical CSE Reference Community composed of 27 communities (Figure 6A). This additional filter based on co-expression metadata could then be layered onto a conventional co-expression network (based on any treatment, developmental stage, or tissue type selected by the researcher), and thus provide a more detailed and nuanced view of the innate relationships between the genes, when stress/treatment/tissue/genotype effects have been nullified.

**Figure 6.**
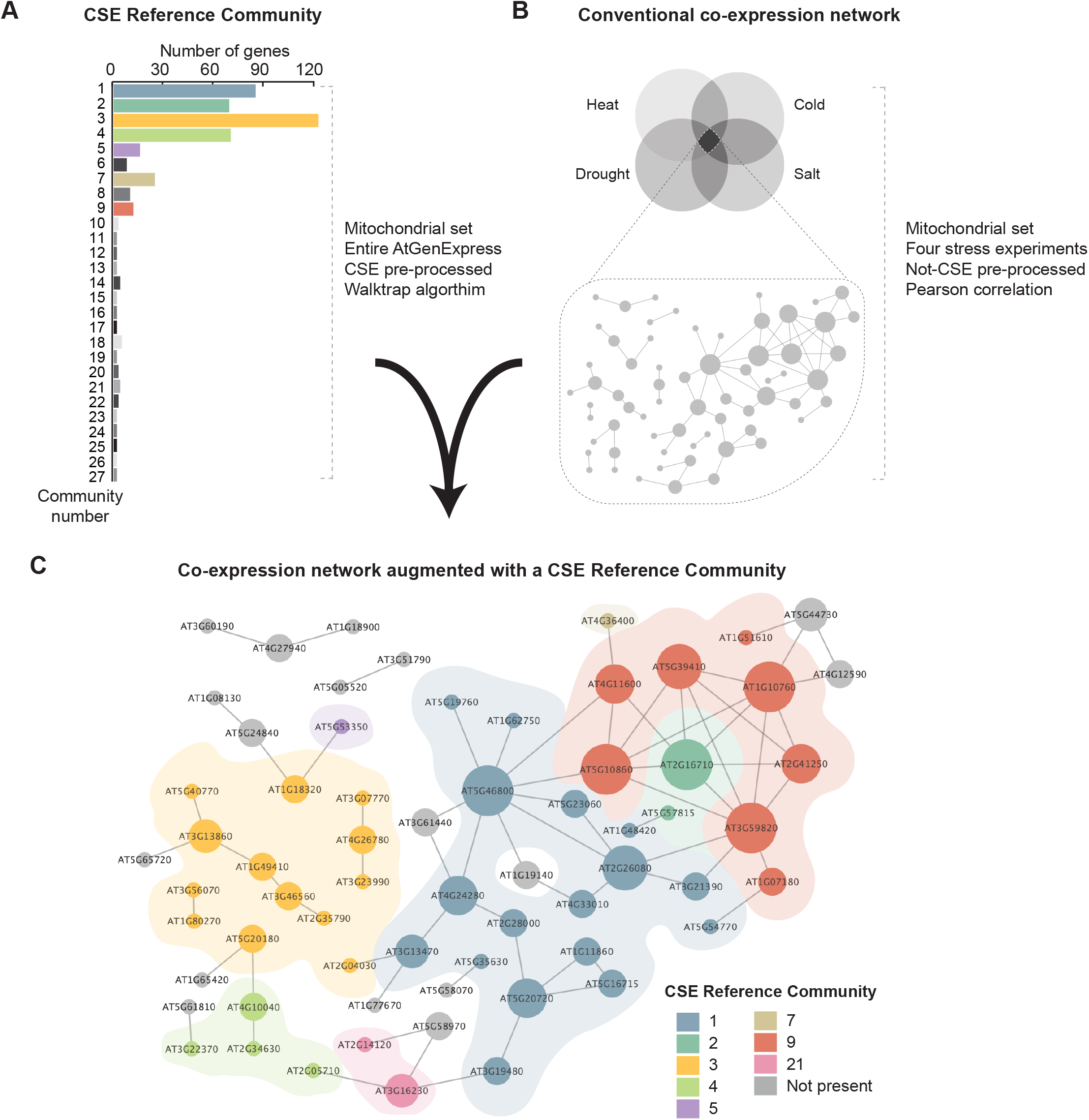
Synthesis of a conventional co-expression network of Arabidopsis shoots common to four stresses with a CSE Reference Community. **(A)** A CSE Reference Community was generated utilising the entire AtGenExpress Expression Atlas, using CSE pre-processed data. This network was divided into 27 primary clusters using a Walktrap community detection algorithm (Pons and Latapy, 2005). **(B)** A core set of stress-responsive genes was isolated from the AtGenExpress stress dataset (Kilian et al., 2007) covering Heat, Drought, Cold and Salt stresses and from this, a network was generated based on Pearson correlation coefficient with no CSE. **(C)** The initial network of non-centralised core stress response generated using Pearson correlation coefficient was cross-referenced with the centralised reference communities; providing deeper insight into the connectivity between genes, independent of outside influences such as stress or tissue type. The diameter of each node is proportional to the number of edges it has to a neighbouring node and node colouration denotes occupation within a given CSE Reference Community.

To illustrate this, we identified a subset of 65 mitochondrial genes that are highly co-expressed in shoot tissues in response to the following four stress treatments: Heat, Cold, Drought, and Salt, using non-CSE pre-processed data (Kilian et al., 2007). As shown in Figure 6B, conventional co-expression analysis (here based on Pearson correlation coefficient) provides an initial network, which illustrates the influence of stress on the relationship between specific stress-responsive genes. When the expression network of the core stress responsive gene was cross-referenced with the CSE Reference Community, the resulting subdivisions revealed unique insights into the functional composition and basal connectivity of this network (Figure 6C). For example, most of genes grouped in Community 1 were associated with photorespiration and thiamine biosynthesis, two metabolic pathways often associated with stress response in plants, and notably in photosynthetic parts (Supplemental Figure 2; Hodges et al., 2016; Rapala-Kozik et al. 2012). Furthermore, Community 3 was overwhelmingly composed of functions associated with translation (e.g. ribosomal protein L36), import (e.g. TOM6, TIM9, and the TIM-family protein AT1G18320), and assembly (e.g. HSP60-3A, HSP6, Hsp89.1, CR88, and MGE2). Interestingly, a number of the genes in this shoot core stress set were also present in a corresponding network prepared from root data (denoted with a black outline in Figure 6B). Of these shared genes, 2/3^rd^ are found in Community 3, which again emphasises their importance. Therefore, we propose that viewing traditional co-expression networks through a prism of a CSE Reference Community can rapidly reveal hidden degrees of connectivity between genes and could have far-reaching applications in the field of transcriptomics, regardless of organisms, treatments or even pathologies.

## Discussion

In light of the burgeoning output of next generation sequencing projects performed on any species, under different developmental or clinical conditions, the statistical power and complexity of these networks will undoubtedly increase, while their biological relevance will be fiercely challenged. Therefore, it is essential that current methodologies be refined to keep apace of this progress and utilise these resources to generate more accurate and informative gene networks to answer hypothesis-driven questions. With the present study, we proposed an alternative method to conventional batch corrections and demonstrated that the implementation of CSE (performed simultaneously per gene and per sub-experiment) and used in isolation or coupled to traditional correlation approaches, can provide additional biological relevance to conventional co-expression networks.

Arguably, there is not a unique defining co-expression network, since the degree of co-expression clearly depends on the context considered, e.g. tissue, growth condition or developmental stage. Nonetheless, we believe there is utility in generating a core network, where the edges corresponds to essential interactions and pathways that are commonly present. Furthermore, the predicted number of edges in a GCN is determined by user defined inclusion rules, e.g. an edge is predicted if the correlation is significant and/or has a value greater than a given threshold. From a biological point of view, these inclusion criteria are problematic since the number of edges depends on the number of samples (the more samples, the lower the p-values and thus the more edges) and which method is used to quantify the co-expression. For example, GCNs using CSE will on average estimate less extreme correlations than GCNs not using CSE, although they may share several edges (see Supp. Fig. 3). We argue that a sensible alternative approach is to control the sparsity of the network and to consider the predicted edges simply as the most pronounced co-expressions.

The predicted core network depends on the coverage of included samples, which necessitates extensive sampling; covering different tissue types, developmental stages, and stresses. A challenge of sampling broadly is the difficulty of combining samples from contextually different experiments, with core gene co-expression being obscured by treatment-associated co-expression. One solution would be to split the experimental data into subsets where each subset consists of data from similar experiments, and predict a separate network for each dataset, and finally estimate the core network with a consensus network. However, this approach suffers from some shortcomings; it may be difficult to define the subsets, there may be relatively few samples within the subsets and it is unclear how to derive the consensus network. The proposed pre-processing method CSE, which can be combined with any GCN method, defines the subsets (i.e. the sub-experiments) conservatively and mechanically, where each sub-experiment consists of biological replicates, and removes all treatment effects including batch effects, allowing for a direct estimation of the core network based on all available samples. A drawback with the CSE approach is that it will reduce the signal-to-noise ratio. For the considered Arabidopsis data, with 887 samples, this seems to be minor problem, but for relative small data sets it remains an open question whether this could become a hurdle.

Evaluation and validation of GCNs is a challenge, since we have limited information on the “true” relationship that exists between genes. We commonly have experimentally confirmed protein-protein interactions and for some subsets of genes it may be reasonable to assume a relatively high degree of co-expression. We usually lack information on truly non-existing edges. In fact, from a theoretical point of view, we may argue that all pairs of genes are co-expressed to some extent. We propose that the validation should be based on pronounced sub-networks for which we expect to observe more co-expression (i.e. more edges) than expected by chance. This approach allows us to compare different GCNs, all with the same sparsity, and to easily access statistical significance. It should be stressed that the result of the validation depends on the sparsity level and which pronounced sub-networks are used in this validation. In particular, if the number of genes is high it may be recommended to construct a relatively dense network and to include several pronounced sub-networks to ensure high power of the tests.

Here, we used a plant mitochondrial case study, where a series of validation steps established the strength of GCNs built upon data that had been pre-processed with CSE. Plant mitochondria are highly adaptive organelles that can tailor their protein complement to undertake a multitude of specialised roles. Nonetheless, there are a set of canonical functions and associated pathways that are maintained/operated in most tissues, growth conditions, developmental stage, etc. even though such pathways (e.g. respiration, TCA cycle, amino acids catabolism) can of course be differentially regulated to modulate their intensity i.e. regulate the metabolic flux through them. This means that the genes encoding proteins involved in those pathways are functionally correlated even though their respective expression profiles may diverge slightly to satisfy a certain metabolic modularity. Our results show that CSE-based GCNs had significantly more edges within the majority of the considered pronounced sub-networks (i.e. the mETC-complex and its sub-complexes and networks defined by annotation) than GCNs not using CSE (Fig. 3, Fig. 4); which demonstrates that the CSE-based GCNs are efficient at predicting those canonical functions and associated pathways, also referred to as the core network. Furthermore, we showed that CSE, in conjunction with conventional Pearson correlation can be used to fine-tune the prediction of the function for uncharacterised genes (Fig. 5); while a combination with non-centralised data can augment conventional stress analyses with the innate connections underpinning the dynamic system examined (Fig. 6). Indeed, the trade-off of a CSE approach is that the biological precision gained by strengthening a core gene-network results in a loss of information from any stress/treatment/genotype components of the dataset. Despite this, if the focus of a given study is centred on determining the network articulated around specific stress-responsive genes, one can apply a CSE Reference Community onto a conventional “stress” co-expression network. This augments the network with extended biological insights, and provides the user with a resource to better interrogate the biological context of the data. Such context is often hindered by the use of stringent cut-offs and thresholds throughout the gene network establishment. Finally, although based on a plant mitochondrial set to simplify the biological validation of our method, the present study provides an alternative method for interrogating the biological relevance of any gene co-expression network, regardless of organism or biological context.

## Methods

### Dataset generation

To obtain the widest coverage possible of a plant transcriptome, the AtGenExpress expression atlas was utilised. This resource is the result of a multinational consortium that aimed to define an exhaustive transcriptome, covering (i) Arabidopsis developmental stages and tissues types (Schmid et al., 2005), (ii) biotic and abiotic stress treatments (Killian et al., 2007), and (iii) hormone and chemical treatments (Goda et al., 2008). These studies used Affymetrix ATH1 arrays and, where possible, maintained consistent experimental practices between samples so as to optimise comparability. For this study, 887 CEL files from the AtGenExpress set (spanning over 370 unique experimental conditions) were quantile normalised together resulting in the pre-processed dataset. For each unique condition (henceforth referred to as sub-experiment) there were two or three samples, which can be regarded as biological replicates observed under similar conditions, where the conditions were defined with respect to tissue developmental stage and treatment (e.g. a different type of stress). See Supplemental Table 1 (https://www.upsc.se/researchers/4638-olivier-keech-stress-induced-senescence-and-its-subsequent-metabolic-regulations.html#resources).

### Construction of Gene Co-expression Networks

All analysis, if nothing else is said, was conducted with the statistical programming language R version (R 3.5.1) (R Core Team, 2018). The R-code used to construct the GCNs described below are found in our GitHub repository (Kellgren and Rydén, 2019; https://github.com/Tezinha/Gene-Co-expression-Network).

The precision matrices were derived by controlling the fraction of edges in the off-diagonal precision matrix at a user defined level ω. The elements of the precision matrix were derived from a correlation matrix where the elements were set to “one” if the absolute value of the correlations were larger than a cut-off α, and “zero” otherwise. The threshold α was obtained by an iterative procedure controlling the sparsity at the level ω=0.005.

The above approach was used for all analyses with the exception of exception of the analysis resulting in the predicted communities presented in Figure 6 where an alternative bootstrap approach was used. Here samples were randomly chosen with replacement, followed by calculation of the precision matrix as described above. This procedure was repeated 50 times and the resulting precision matrices were combined, generating a matrix with values ranging from 0 to 50. The elements of the precision matrix were derived from the aggregated matrix, where the elements were set to “one” if the values exceeded a cut-off β, and “zero” otherwise. Here β was chosen to control the sparsity ω at 0.005.

Due to computational reasons partial correlation approaches are often carried out on subsets of genes, rather than the whole genome of an organism. An example of this was detailed in Ma et al., (2007), which used a modified GCN approach to carry out partial correlation analysis on batches of ~2000 genes at a time. Aided by iterative random samplings of genes, this study increased their coverage to that of the Affymetrix ATH1 array; resulting in a network composed of 18 625 interactions (edges) and 6760 genes (nodes) (Ma et al., 2007). Ren et al., (2015) expanded on this and proposed an algorithm for constructing GCN with high-dimensional data by implementing asymptotically normal estimation of large GCN, and in doing so, made it realistic to perform GCN at a whole-genome scale (Wang et al., 2016). Unsurprisingly, this approach is enormously computationally taxing, which can prove prohibitive to researchers lacking dedicated servers and advanced computer processing power.

### Evaluation of Gene Co-expression Networks

For any sub-network *A*, with *n*_*A*_ nodes and *k*_*A*_ observed edges, of the predicted core network C, with n nodes and sparsity ω, it possible to test if the sub-network is pronounced (the hull hypothesis) versus that the sub-network is not pronounced (the null hypothesis). Under the null hypothesis *k*_*A*_ is binomial distributed, i.e.

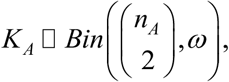

and a binomial test can be used to derive a p-value. Here the R-function “binom.test” (R 3.5.1) was used to derive the p-values.

It should be stressed that the p-values will depend on the network’s sparsity as well as the size of the sub-network, the larger the pronounced sub-networks are the lower p-values will be expected. Hence, all tough not necessary, having the same sparsity in all networks simplifies the evaluation. Moreover, the more pronounced sub-networks that can be correctly identified the more reliable the evaluation will be.

### Preparing elements of the mitochondrial working model

#### (i) Defining the mitochondrial gene list

The manually curated list of genes encoding proteins targeted to the mitochondria from Chrobok et al., (2016) was used as a basis for a mitochondrial case-study. Matching this list with the AtGenExpress Expression Atlas resulted in a list of 984 mitochondrial genes, which were used for downstream analysis. The samples were taken from different tissues: flower, root, shoot, seedling, leaf, pollen and silique. Mitochondrial genes were categorised with respect to expression patterns, functional proximity and functional categories for downstream validation (Supplemental Table 1).

#### (ii) Defining below-ground and above-ground dominant genes

The mitochondrial genes were classified in two categories with respect to their expression patterns in below-ground tissues (e.g. root) and above-ground tissues (e.g. shoot and leaf). For each gene *i*, the difference between the mean expressions in below-ground tissues, 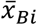 and above-ground tissues, 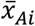 was calculated, i.e.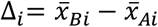. Genes with a difference larger than one standard deviation, i.e. Δ_i_ > *s*_Δ_ were classified as *below-ground dominant genes*, while those with a difference smaller than one standard deviation, i.e. Δ_*i*_ < −*s*_Δ_ were classified as *above-ground dominant genes.* The estimated standard deviation was based on all the Δ-values of genes.

#### (iii) Defining components of Complex I of the mitochondrial electron transport chain

Complex I of the mitochondrial electron transport chain (mETC) was an ideal model to test the effect of functional proximity of the resulting networks, as the identity and molecular arrangement of these constituents have been thoroughly characterised in Arabidopsis using proteomic approaches (Klodmann and Braun., 2010; Peters et al., 2013).

#### (iv) MapMan annotations

Using the newly updated functional annotations established for the MapMan platform (MapMan X4 Release 1.0, 2018; Usadel et al., 2009), each gene of the mitochondrial set was assigned to one of 29 functional categories.

### Preparing a Reference Community Set

The *Walktrap community detection algorithm* runs short random walks and merges separate communities in a bottom-up manner to produce clusters, and was applied to the derived networks to identify *gene communities*, i.e. sets of genes with a high degree of predicted intra-gene-gene interactions. The function “walktrap.community” with default settings in the R package igraph (Csárdi and Nepusz, 2006) was used to conduct the analyses. Here, gene communities were predicted based on a network obtained using centralized data from all experiments, Pearson correlation and a precision matrix derived using the absolute value of the correlations. The result was a CSE Reference Community composed of 27 clusters.

### Combining results obtained using centralized and non-centralized data

We claim that gene communities should be estimated based on networks derived using all the available centralized data, while networks based on non-centralized data describe how genes are affected by an external factor, e.g. stress induced by heat, cold, salt or drought. Combining the two type of networks allowed us to study how gene communities were affected by stress. The combined analysis was made as follows. First the communities were predicted as described above, resulting in the *community network*. Secondly, for each of the considered stresses, samples exposed to the stress were selected (heat n=16, cold n=24, salt n=24, and drought n=28). A precision matrix was calculated using non-centralized data, Pearson correlation, and non-bootstrap approach with a cut-off=0.82. The sum of the four stress-related precision matrices was calculated and edges with an aggregated score equal to 4 were set to “one” in the combined precision matrix (i.e. the *stress network*) and regarded as gene-gene interaction caused by a general stress response.

The community and stress networks were combined. Communities enriched with respect to general stress were identified similarly as described above. An enrichment analysis with respect to functional categories was made for each of the enriched communities.

## Declarations

### Ethics approval and consent to participate

Not applicable

### Consent for publication

Not applicable

### Availability of data and material

- The datasets analysed during the current study are available under the AtGenExpress expression atlas, which is the result of a multinational consortium that aimed to define an exhaustive transcriptome, covering i) Arabidopsis developmental stages and tissues types (Schmid et al., 2005), ii) biotic and abiotic stress treatments (Killian et al., 2007), and iii) hormone and chemical treatments (Goda et al., 2008).
- All data generated during this study are included in this published article and its supplementary information files.

## Author’s contribution

SL, TK and RB performed analyses. SL prepared the figures and drafted the manuscript, PR and OK conceptualized the project and edited the manuscript. All authors read and approved the manuscript.

---

**Supplemental Figure S1. Schematic representations of the conclusions that can be drawn from different correlation analysis approaches of gene expression data.** Five genes were simulated to illustrate a network in the following way; Gene A expression affects Gene B expression, Gene C expression affects the expression of Gene D and Gene E. The gene’s expression values are regarded as functions of a normally distributed random variable, with a mean μ=0, and a standard deviation σ=0.5. The expression of two of the genes, Gene A and Gene C are also affected by an external stress treatment, which can be seen as a categorical variable with two levels. Level one represent no external influences and the variable takes a value of zero, at level two the gene is influenced by an external factor and the categorical variable takes the value ten. Gene B expression is affected by the expression of Gene A, so for each Gene B value a Gene A value multiplied by a constant β=0.5 is added. In the same way, Gene D and Gene E is simulated but with the exception that they are affected by Gene C. For each of the scenarios 100 expression values were simulated for each gene. To compare Pearson’s correlation against partial correlation the relative correlation, i.e. the most correlated edge, was set as a baseline and received a correlation value of 1. This was done for each setup. In the first column the true network is represented and if it is affected by the external factor. In column 2 to 5 the strength of the relative correlations is represented by the thickness of the line. **(A)** The network is not affected by any external factor and all four methods have the correct edges among the top three candidates. There is no difference between non-centralised and centralised data which is as expected when there is no external factor to remove with CSE. **(B)** The stress treatment is affecting gene C expression, which has an effect on the non-centralised networks. Pearson correlation gives a false positive among the top three candidates, the partial correlation networks gives the correct top three candidates but the edge between Gene A and B is weak. When we preform CSE both networks give the correct top three edges. **(C)** In this case, the stress treatment is affecting the expression of both Gene A and C, which leads to false positives with both methods. By carrying out CSE, the stress treatment, is removed and both Pearson and partial correlation output the correct top three edges.

**Supplemental Figure S2. Synthesis of a conventional co-expression network of Arabidopsis shoots common to four stresses with a CSE Reference Community Set.** A core set of stress-responsive genes isolated from non-centralised AtGenExpress stress dataset (Kilian et al., 2007) covering Heat, Drought, Cold and Salt stresses, cross-referenced with the CSE Reference Community.

**Supplemental Figure S3. Correlation between the 985 mitochondrion related genes were estimated using Pearson correlation without centralization (Non-Centralized data) and Pearson correlation with CSE preprocessing (CSE preprocessed data).** For each approach 484,620 correlations were estimated and the 0.5 % (2423) gene correlations with the highest absolute value were used to predict edges in the corresponding gene co-expression network. (A) Estimated density functions over all estimated correlations for non-centralized data (green) and CSE preprocessed data (red). The black line shows the density for correlations estimated on simulated noise. (B) The estimated correlations for the two approaches plotted against each other. Edges shared by both approaches are marked blue (620 (25.6 %) of the edges were shared), unique edges for the CSE preprocessing network are marked red, and unique edges for the Non-centralized network are marked green.

**Supplemental Table 1.** List of 984 genes encoding proteins targeted to the mitochondrion, referenced with the AtGenExpress Expression Atlas (Schmid et al., 2005, Kilian et al., 2007 and Goda et al., 2008). Note that dues to its large size (ca. 250 MB), the file is available at: https://www.upsc.se/researchers/4638-olivier-keech-stress-induced-senescence-and-its-subsequent-metabolic-regulations.html#resources

**Supplemental Table 2. i)** Non-CSE Pearson correlation; **ii)** Non-CSE Partial correlation; **iii)** CSE Pearson correlation; **iv)** CSE Partial correlation.

**Supplemental Table 3. Statistics supporting Figure 3.** Table of the expected, observed, ratios, and associated P-values. This is carried out for interactions within Complex I, within and between the 5 Complexes of the mETC, and between the mETC and the rest of the mitochondrion.

**Supplemental Table 4.** List of source and target genes comprising genes encoding proteins targeted to the mitochondrion, with unknown functions (as per MapMan X4 annotations) and their edges with known mitochondrial genes.

**Supplemental Table 5.** Table of the 27 communities generated using the Walktrap algorithm on the whole AtGenExpress Set that has been centralised

## References

Banf M, Rhee SY (2017) Computational inference of gene regulatory networks: Approaches, limitations and opportunities. BBA Gene reg mech 1860(1):41–52. Epub 2016/09/20.

Barkan A, Small I (2014) Pentatricopeptide repeat proteins in plants. Annu Rev Plant Biol 65: 415–442

Carrera J, Rodrigo G, Jaramillo A, Elena SF (2009) Reverse-engineering the *Arabidopsis thaliana* transcriptional network under changing environmental conditions. Genome Biol 10:R96

Castro DM, de Veaux NR, Miraldi ER, Bonneau R (2019) Multi-study inference of regulatory networks for more accurate models of gene regulation. PLoS comp biol 15(1):e1006591.

Chai LE, Loh SK, Low ST, Mohamad MS, Deris S, Zakaria Z (2014) A review on the computational approaches for gene regulatory network construction. Comp biol med. 48:55–65. Epub 2014/03/19.

Cheng CY, Krishnakumar V, Chan A, Schobel S, Town CD (2017) Araport11: a complete reannotation of the *Arabidopsis thaliana* reference genome. Plant J 89:789–804.

Chen C, Grennan K, Badner J, Zhang D, Gershon E, et al. (2011) Removing Batch Effects in Analysis of Expression Microarray Data: An Evaluation of Six Batch Adjustment Methods. PLoS ONE 6(2): e17238. doi:10.1371/journal.pone.0017238

Chrobok D, Law SR, Brouwer B, Lindén P, Ziolkowska A, Liebsch D, Narsai R, Szal B, Moritz T, Rouhier N, Whelan J, Gardeström P, Keech O (2016) Dissecting the metabolic role of mitochondria during developmental leaf senescence. Plant Physiol 172: 2132–2153

Csárdi G, Nepusz T (2006) The igraph software package for complex network research, InterJournal, Complex Systems 1695

Doniwa Y, Ueda M, Ueta M, Wada A, Kadowaki K, Tsutsumi N (2010) The involvement of a PPR protein of the P subfamily in partial RNA editing of an Arabidopsis mitochondrial transcript. Gene; 454: 39–46

Emmert-Streib F, Dehmer M, Haibe-Kains B (2014) Gene regulatory networks and their applications: understanding biological and medical problems in terms of networks. Front Cell Dev Biol 2:38

Goda H, Sasaki E, Akiyama K, Maruyama-Nakashita A, Nakabayashi K, Li W, Ogawa M, Yamauchi Y, Preston J, Aoki K, Kiba T, Takatsuto S, Fujioka S, Asami T, Nakano T, Kato H, Mizuno T, Sakakibara H, Yamaguchi S, Nambara E, Kamiya Y, Takahashi H, Hirai MY, Sakurai T, Shinozaki K, Saito K, Yoshida S, Shimada Y (2008) The AtGenExpress hormone and chemical treatment data set: experimental design, data evaluation, model data analysis and data access. Plant J 55: 526–542

Ghifari AS, Gill-Hille M, Murcha MW (2018) Plant mitochondrial protein import: the ins and outs. Biochem J 475 (13) 2191–2208

Haïli N, Planchard N, Arnal N, Quadrado M, Vrielynck N, Dahan J, des Francs-Small CC, Mireau H (2016) The MTL1 pentatricopeptide repeat protein is required for both translation and splicing of the mitochondrial NADH DEHYDROGENASE SUBUNIT7 mRNA in Arabidopsis. Plant Phys 170, 354–366.

Han D, Oh J, Kim K, Lim H, Kim Y (2007) Crystal structure of YrrB: a TPR protein with an unusual peptide-binding site. Biochem Biophys Res Commun 360: 784–790

Hodges M, Dellero Y, Keech O, Betti M, Raghavendra AS, Sage R, Zhu XG, Allen DK, Weber AP (2016) Perspectives for a better understanding of the metabolic integration of photorespiration within a complex plant primary metabolism network. J Exp Bot 67: 3015–3026

Hu C, Lin, SY, Chi, WT, Charng YY (2012) Recent gene duplication and subfunctionalization produced a mitochondrial GrpE, the nucleotide exchange factor of the Hsp70 complex, specialized in thermotolerance to chronic heat stress in Arabidopsis. Plant Physiol 158, 747–758

Kilian J, Whitehead D, Horak J, Wanke D, Weinl S, Batistic O, D’Angelo C, Bornberg-Bauer E, Kudla J, Harter K (2007) The AtGenExpress global stress expression data set: protocols, evaluation and model data analysis of UV-B light, drought and cold stress responses. Plant J 50: 347–363

Kim JH, Han JM, Kim S (2014) Protein–Protein Interactions and Multi-component Complexes of Aminoacyl-tRNA Synthetases. In: Kim S. (eds) Aminoacyl-tRNA Synthetases in Biology and Medicine. Topics in Current Chemistry, vol 344. Springer, Dordrecht

Klodmann J, Sunderhaus S, Nimtz M, Jänsch L, Braun HP (2010) Internal architecture of mitochondrial complex I from *Arabidopsis thaliana*. Plant Cell 22: 797–810

Liesecke F, Daudu D, Dugé de Bernonville R, Besseau S, Clastre M, Courdavault V, de Craene JO, Crèche J, Giglioli-Guivarc’h N, Glévarec G, Pichon O … Dugé de Bernonville T (2018) Ranking genome-wide correlation measurements improves microarray and RNA-seq based global and targeted co-expression networks. Sci Rep 8(1):10885.

Lindén P, Keech O, Stenlund H, Gardeström P, Moritz T (2016) Reduced mitochondrial malate dehydrogenase activity has a strong effect on photorespiratory metabolism as revealed by 13C labelling. J Exp Bot 67(10): 3123–35

Ma S, Gong Q, Bohnert HJ (2007) An Arabidopsis gene network based on the graphical Gaussian model. Genome Res 17:1614–1625

Ma S, Bohnert HJ, Dinesh-Kumar SP (2015) AtGGM2014, an Arabidopsis gene co-expression network for functional studies. Sci China Life Sci 58:3

Manavski N, Guyon V, Meurer J, Wienand U, Brettschneider R (2012) An essential pentatricopeptide repeat protein facilitates 5′maturation and translation initiation of rps3 mRNA in maize mitochondria. Plant Cell 24:3087–3105.

Nygaard V, Rødland E. A, Hovig E (2016). Methods that remove batch effects while retaining group differences may lead to exaggerated confidence in downstream analyses. Biostatistics 17 29–39

Peters K, Belt K, Braun HP (2013) 3D gel map of arabidopsis complex I. Front Plant Sci 4, 153.

Pons P, Latapy M (2005) Computing communities in large networks using random walks. Comp Info Sci; 3733:284–93.

Qian X, Dougherty ER (2013) Validation of gene regulatory network inference based on controllability. Frontiers in genetics. 2013;4:272. Epub 2014/01/01.

R Core Team (2018). R: A language and environment for statistical computing. R Foundation for Statistical Computing, Vienna, Austria. URL https://www.R-project.org/.

Rapala-Kozik M, Wolak N, Kujda M, Banas AK (2012) The upregulation of thiamine (vitamin B1) biosynthesis in *Arabidopsis thaliana* seedlings under salt and osmotic stress conditions is mediated by abscisic acid at the early stages of this stress response. BMC Plant Biol 12: 2–2

Ren Z, Sun T, Zhang C-H, Zhou HH (2015) Asymptotic normality and optimalities in estimation of large Gaussian graphical models. Ann Statist 43(3):991–1026.

Schmid M, Davison TS, Henz SR, Pape UJ, Demar M, Vingron M, Scholkopf B, Weigel D, Lohmann JU (2005) A gene expression map of *Arabidopsis thaliana* development. Nat Genet 37: 501–506

Schmitz-Linneweber C, Small I (2008) Pentatricopeptide repeat proteins: a socket set for organelle gene expression. Trends Plant Sci 13: 663–70

Shannon P, Markiel A, Ozier O, Baliga N, Wang J, Ramage D, Amin N, Schwikowski B, Ideker T (2003) Cytoscape: a software environment for integrated models of biomolecular interaction networks. Genome Res 13:2498–2504.

Tomaz T, Bagard M, Pracharoenwattana I, Lindén P, Lee CP, Carroll AJ, Stroher E, Smith SM, Gardeström P, Millar AH (2010). Mitochondrial Malate Dehydrogenase Lowers Leaf Respiration and Alters Photorespiration and Plant Growth in Arabidopsis. Plant Phys 154: 1143–1157

Usadel B, Obayashi T, Mutwil M, Giorgi FM, Bassel GW, Tanimoto M, Chow A, Steinhauser D, Persson S, Provart NJ (2009) Co-expression tools for plant biology: Opportunities for hypothesis generation and caveats. Plant Cell Environ 32: 1633–1651

Wang T, Ren Z, Ding Y, Fang Z, Sun Z, MacDonald ML, Sweet RA, Wang J, Chen W (2016) FastGGM: An Efficient Algorithm for the Inference of Gaussian Graphical Model in Biological Networks. PLoS comp biol 12(2):e1004755

Wille A, Zimmermann P, Vranová E, Fürholz A, Laule O, Bleuler S, Hennig L, Prelić A, von Rohr P, Thiele L, Zitzler E, Gruissem W, Bühlmann P (2004) Sparse graphical Gaussian modeling of the isoprenoid gene network in *Arabidopsis thaliana*. Genome Biol 5:R92

