## Supplemental Figures for "Enhancing the biological relevance of Gene Co-expression Networks: A plant mitochondrial case study"

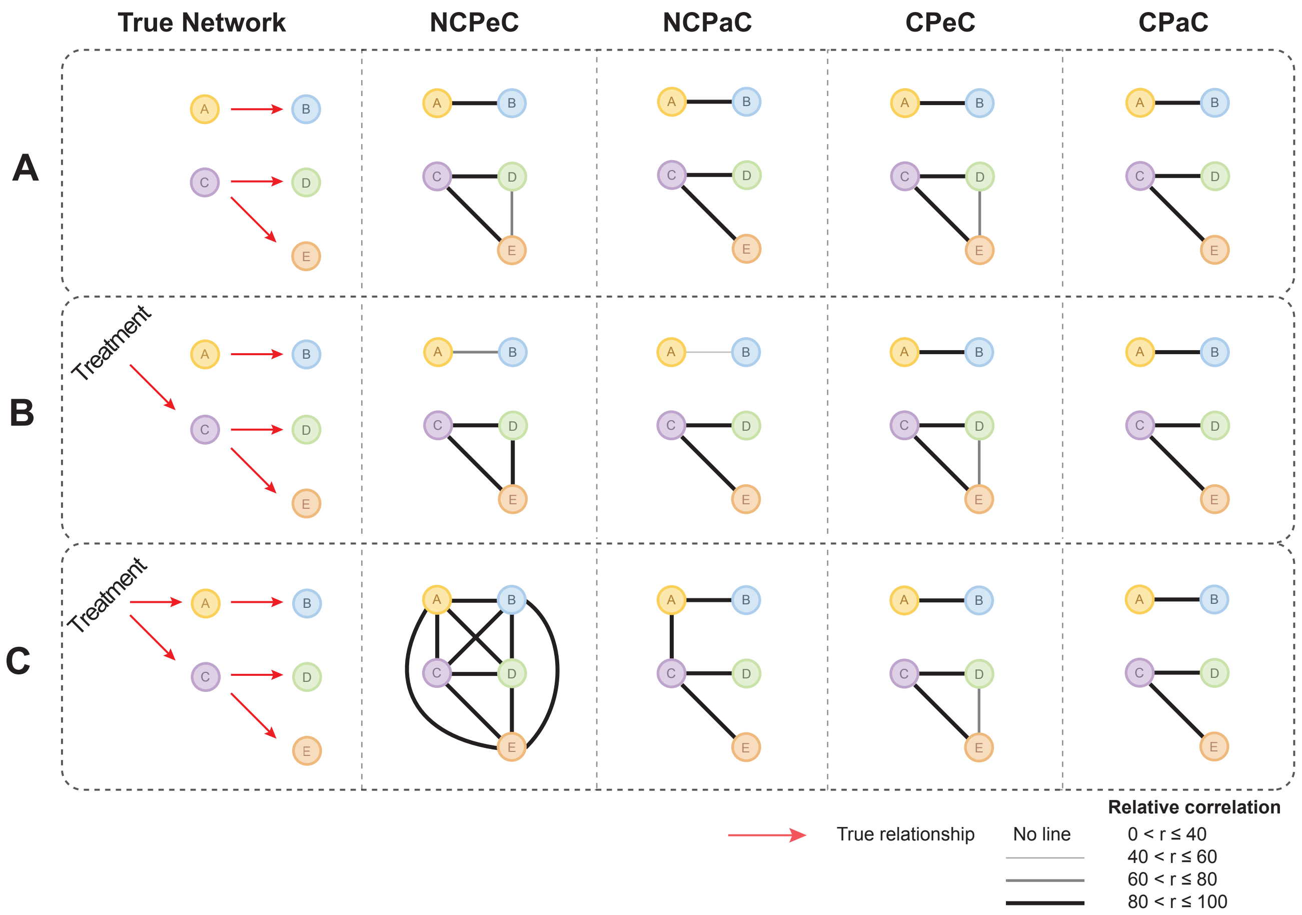


**Supplemental Figure 1. Schematic representations of the conclusions that can be drawn from different correlation analysis approaches of gene expression data.** Five genes were simulated to illustrate a network in the following way; Gene A expression affects Gene B expression, Gene C expression affects the expression of Gene D and Gene E. The gene's expression values are regarded as functions of a normally distributed random variable, with a mean $\mu=0$, and a standard deviation $\sigma=0.5$. The expression of two of the genes, Gene A and Gene C are also affected by an external stress treatment, which can be seen as a categorical variable with two levels. Level one represent no external influences and the variable takes a value of zero, at level two the gene is influenced by an external factor and the categorical variable takes the value ten. Gene B expression is affected by the expression of Gene A, so for each Gene B value a Gene A value multiplied by a constant $\beta=0.5$ is added. In the same way, Gene D and Gene E is simulated but with the exception that they are affected by Gene C. For each of the scenarios 100 expression values were simulated for each gene. To compare Pearson's correlation against partial correlation the relative correlation, i.e. the most correlated edge, was set as a baseline and received a correlation value of 1. This was done for each setup. In the first column the true network is represented and if it is affected by the external factor. In column 2 to 5 the strength of the relative correlations is represented by the thickness of the line. **(A)** The network is not affected by any external factor and all four methods have the correct edges among the top three candidates. There is no difference between non-centralised and centralised data which is as expected when there is no external factor to remove with CSE. **(B)** The stress treatment is affecting gene C expression, which has an effect on the non-centralised networks. Pearson correlation gives a false positive among the top three candidates, the partial correlation networks gives the correct top three candidates but the edge between Gene A and B is weak. When we preform CSE both networks give the correct top three edges. **(C)** In this case, the stress treatment is affecting the expression of both Gene A and C, which leads to false positives with both methods. By carrying out CSE, the stress treatment, is removed and both Pearson and partial correlation output the correct top three edges.

| Reference Community | Locus ID | Symbol | ARAPORT11 Description | Functional Enrichment |
|---|---|---|---|---|
| 1 | AT2G28000 | CPN6-A | chaperonin-60alpha | |
| 1 | AT3G13470 | CPN6-BETA2 | TCP-1/cpn60 chaperonin family protein | |
| 1 | AT5G20720 | CPN2 | chaperonin 20 | |
| 1 | AT4G33010 | GLDP1 | glycine decarboxylase P-protein 1 | |
| 1 | AT2G26080 | GLDP | glycine decarboxylase P-protein 2 | **Photorespiration** |
| 1 | AT1G11860 | - | Glycine cleavage T-protein family | |
| 1 | AT5G46800 | BOU | Mitochondrial substrate carrier family protein | |
| 1 | AT5G35630 | GS2 | glutamine synthetase 2 | |
| 1 | AT3G19480 | 3-PGDH | D-3-phosphoglycerate dehydrogenase | |
| 1 | AT5G19760 | - | Mitochondrial substrate carrier family protein | |
| 1 | AT3G21390 | - | Mitochondrial substrate carrier family protein | **Thiamin biosynthesis** |
| 1 | AT5G54770 | THI1 | thiazole biosynthetic enzyme | |
| 1 | AT5G16715 | EMB2247 | protein EMBRYO DEFECTIVE 2247 | |
| 1 | AT1G62750 | SCO1 | Translation elongation factor EFG/EF2 protein | |
| 1 | AT4G24280 | cpHsc7--1 | chloroplast heat shock protein 70-1 | |
| 1 | AT5G23060 | CaS | calcium sensing receptor | |
| 1 | AT1G48420 | D-CDES | D-cysteine desulfhydrase | |
| 2 | AT5G57815 | - | Cytochrome c oxidase, subunit Vib family protein | |
| 2 | AT2G16710 | - | Iron-sulfur cluster biosynthesis family protein | |
| 3 | AT5G20180 | - | Ribosomal protein L36 | |
| 3 | AT3G56070 | ROC2 | rotamase cyclophilin 2 | |
| 3 | AT5G40770 | PHB3 | prohibitin 3 | |
| 3 | AT1G49410 | TOM6 | translocase of the outer mitochondrial membrane 6 | |
| 3 | AT1G18320 | - | Mitochondrial import Tim17/Tim22/Tim23 family protein | |
| 3 | AT3G46560 | TIM9 | Tim10/DDP family zinc finger protein | |
| 3 | AT3G13860 | HSP60-3A | heat shock protein 60-3A | **Protein synthesis, import and assembly** |
| 3 | AT3G23990 | HSP6 | heat shock protein 60 | |
| 3 | AT3G07770 | Hsp89.1 | HEAT SHOCK PROTEIN 89.1 | |
| 3 | AT2G04030 | CR88 | Chaperone protein htpG family protein | |
| 3 | AT1G80270 | PPR596 | PENTATRICOPEPTIDE REPEAT 596 | |
| 3 | AT4G26780 | MGE2 | Co-chaperone GrpE family protein | |
| 3 | AT2G35790 | - | transmembrane protein | |

| Reference Community | Locus ID | Symbol | ARAPORT11 Description |
|---|---|---|---|
| 4 | AT2G05710 | ACO3 | aconitase 3 |
| 4 | AT4G10040 | CYTC-2 | cytochrome c-2 |
| 4 | AT3G22370 | AOX1A | alternative oxidase 1A |
| 4 | AT2G34630 | GPS1 | geranyl diphosphate synthase 1 |
| 5 | AT5G53350 | CLPX | CLP protease regulatory subunit X |
| 7 | AT4G36400 | D2HGDH | FAD-linked oxidases family protein |
| 9 | AT1G07180 | NDA1 | alternative NAD(P)H dehydrogenase 1 |
| 9 | AT1G10760 | SEX1 | Pyruvate phosphate dikinase |
| 9 | AT4G11600 | GPX6 | glutathione peroxidase 6 |
| 9 | AT1G51610 | - | Cation efflux family protein |
| 9 | AT5G39410 | - | Saccharopine dehydrogenase |
| 9 | AT5G10860 | CBSX3 | Cystathionine beta-synthase family protein |
| 9 | AT3G59820 | LETM1 | LETM1-like protein |
| 9 | AT2G41250 | - | Haloacid dehalogenase-like hydrolase superfamily protein |
| 21 | AT2G14120 | DRP3B | dynamin related protein |
| 21 | AT3G16230 | - | Putative eukaryotic LigT |
| NP | AT5G58970 | UCP2 | uncoupling protein 2 |
| NP | AT3G51790 | G1 | transmembrane protein G1P-related 1 |
| NP | AT5G65720 | NFS1 | nitrogen fixation S (NIFS)-like 1 |
| NP | AT1G19140 | COQ9 | ubiquinone biosynthesis COQ9-like protein |
| NP | AT1G08130 | LIG1 | DNA ligase 1 |
| NP | AT5G24840 | TRM8A | tRNA (guanine-N-7) methyltransferase |
| NP | AT3G60190 | DL1E | DYNAMIN-like 1E |
| NP | AT5G05520 | - | Outer membrane OMP85 family protein |
| NP | AT5G61810 | APC1 | Mitochondrial substrate carrier family protein |
| NP | AT4G27940 | MTM1 | manganese tracking factor for mitochondrial SOD2 |
| NP | AT1G65420 | NPQ7 | antigen receptor-like protein (DUF565) |
| NP | AT1G77670 | - | Pyridoxal phosphate dependent transferases superfamily protein |
| NP | AT5G58070 | TIL | temperature-induced lipocalin |
| NP | AT1G18900 | - | Pentatricopeptide repeat superfamily protein |
| NP | AT5G44730 | - | Haloacid dehalogenase-like hydrolase superfamily protein |
| NP | AT4G12590 | - | ER membrane protein complex subunit-like protein |
| NP | AT3G61440 | CYSC1 | cysteine synthase C1 |

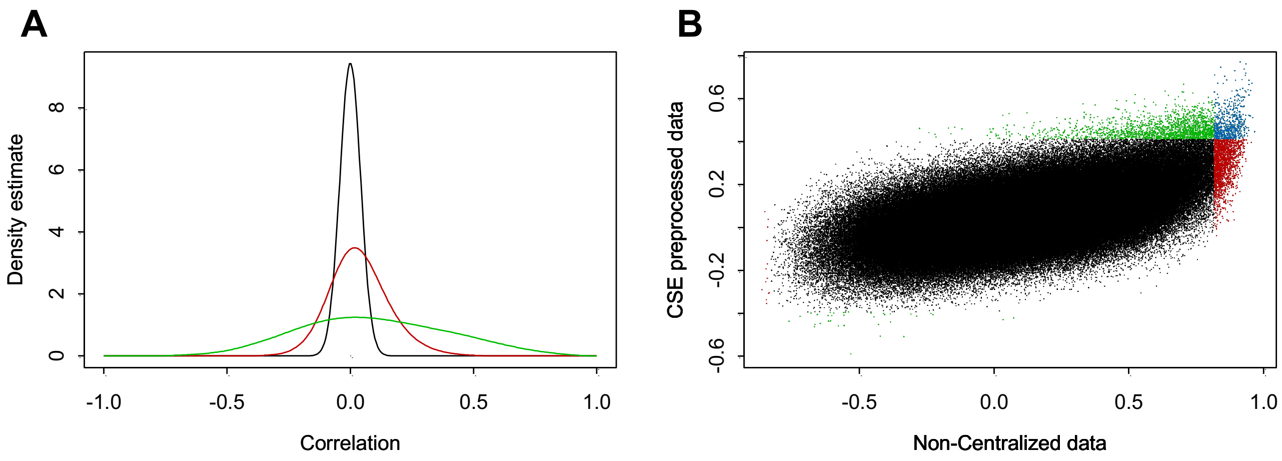


**Supplemental Figure 3. Correlation between the 985 mitochondrion related genes were estimated using Pearson correlation without centralization (Non-Centralized data) and Pearson correlation with CSE preprocessing (CSE preprocessed data)**. For each approach 484,620 correlations were estimated and the 0.5 % (2423) gene correlations with the highest absolute value were used to predict edges in the corresponding gene co-expression network. **(A)** Estimated density functions over all estimated correlations for non-centralized data (green) and CSE preprocessed data (red). The black line shows the density for correlations estimated on simulated noise. **(B)** The estimated correlations for the two approaches plotted against each other. Edges shared by both approaches are marked blue (620 (25.6 %) of the edges were shared), unique edges for the CSE preprocessing network are marked red, and unique edges for the Non-centralized network are marked green.
